# Transcriptional subtypes, anatomical organization, and sexual dimorphism of sensory vagus neurons in *Danionella cerebrum*

**DOI:** 10.64898/2026.09.01.748112

**Authors:** Emily A. Bayer, Alexander F. Schier

## Abstract

Visceral sensory neurons sense and modulate the brain, behavior, and the internal organs. However, these neurons have been difficult to study comprehensively due to their projections deep within the body. Here, we establish the transparent miniature fish *Danionella cerebrum* as a model for studying the sensory vagus nerve in an adult vertebrate. By generating a transgenic line to label *D. cerebrum* cranial sensory ganglia, we were able to both anatomically characterize and transcriptionally profile the sensory vagus at single-cell resolution. Anatomically, we find that the vagal ganglia have a somatotopic layout. Transcriptionally, the sensory vagus is comprised of diverse sensory subtypes conserved with other vertebrates, including nutrient-sensing, mechanoreceptive, nociceptive, and thermosensitive subtypes, as well as polymodal combinations. Visualizing subtype marker genes identified somatotopic-specific sensory subtypes. Notably, the *D. cerebrum* sensory vagus is not static across development: it becomes anatomically sexually dimorphic during sexual maturation and undergoes continuous adult neurogenesis. The molecular and anatomical atlas of the sensory vagus lays the foundation for functional studies of body-brain communication in *D. cerebrum*.

## INTRODUCTION

One fundamental function of the brain is to generate behavioral responses to sensory stimuli. In addition to information from the outside world, the brain constantly receives sensory information from inside of the body. Visceral sensory neurons respond to a variety of stimuli to transmit information from organs including the gut, lung, and heart^1–5^. In turn, this information can be used to regulate organ function and impact overall animal behavior^6^.

The vagus nerve, cranial sensory nerve X, is a major source of visceral sensory input, and contains mostly sensory but also motor components ^7,8^. In mammals, the cell bodies of the vagus nerve are located in two ganglia (jugular and nodose) at the base of the skull, which then project to the internal organs^9^. The nodose ganglion is placode-derived, while the jugular ganglion is neural crest-derived^10^. Increasingly detailed information is available about the molecular subtypes of the sensory vagus and the representation of vagus inputs in the brain. Vagal sensory neurons encode information across multiple dimensions, including sensory modality, target organ, and tissue layer^1^. While some sensory modalities such as stretch sensitivity are broadly represented across the ganglion, the sensory vagus also contains extremely rare cell types: some throat and pulmonary sensory neuron types make up under 1% of the mouse sensory vagus^11,12^. To understand the sensations that the brain receives from inside of the body, it is thus important to comprehensively characterize the cell types and their projections in the sensory vagus.

Teleost fish also possess an elaborate vagal nerve with the ability to sense a variety of stimuli. As in mammals, the teleost vagus is predominantly sensory^13^. For example, the gill arches are extensively innervated, which enables fish to sense and respond to internal stimuli such as heart rate, pharyngeal dilation, and respiratory cycle (reviewed in^14^). The gills serve as the primary sense organs to assess water quality, using chemical and mechanical nociceptors to respond to noxious stimuli, in addition to baroreceptors and chemoreceptors that are associated with vagus afferent fibers^14^. The sensory vagus also extensively innervates the most posterior branchial arch, which includes the pharyngeal jaws and teeth^15^.

In addition to these gill-related functions, the zebrafish placodal vagus (nodose) ganglion projects posteriorly to the visceral organs^15,16^. Functional studies in zebrafish have demonstrated that the vagus nerve transmits stimuli such as distension, nutrient content, and irritants from the gut^17,18^, and vagal sensory neuron activity is correlated with both bradycardia and tachycardia^19^. However, while larval zebrafish is a powerful model for neuroscience, vagal outgrowth and organ innervation are gradual and still immature during larval stages^20^. Thus, it would be advantageous to identify an experimental system that remains accessible through circuit maturation and adulthood.

The miniature teleost *Danionella cerebrum* is a promising emerging model in neuroscience owing to its small transparent body, and smallest brain of any known vertebrate (∼650,000 neurons^21^ compared with ∼10 million in an adult zebrafish^22^). *D. cerebrum* has a rich behavioral repertoire, and its brain is optically accessible in adults^21,23–27^. In addition, its body is transparent, allowing visualization of all the internal organs (**Figure 1A**). These features position *D. cerebrum* as a uniquely powerful system in which to study body-brain signaling and visceral sensation.

**Figure 1:**
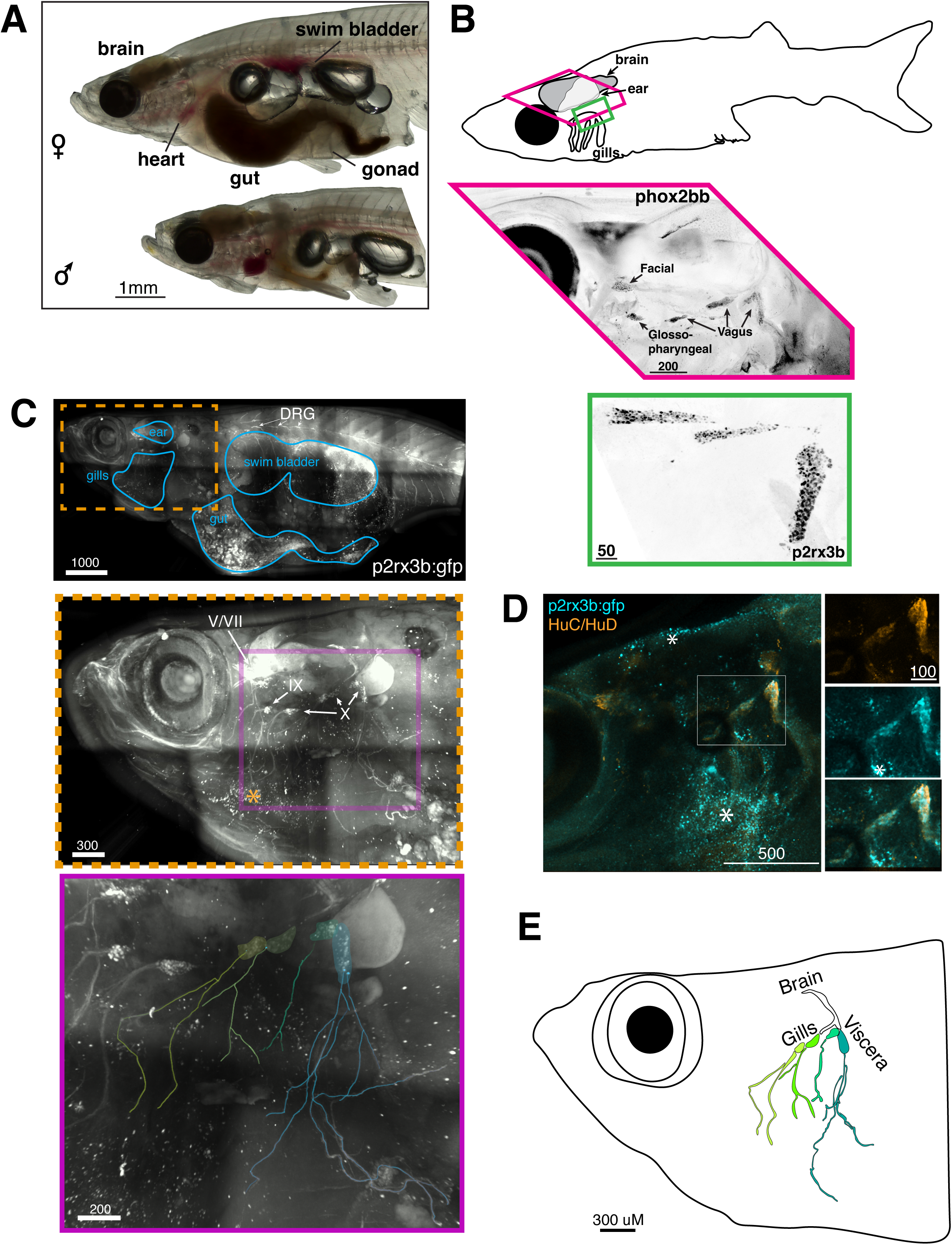
Adult transparency of *D. cerebrum* enables visualization of brain-body connectivity. **A. *D. cerebrum* visceral organs are visible in adult animals.** Bright field images of adult male and female animals (5 months post-fertilization) with major organs labeled. **B. *phox2bb* and *p2rx3b* are expressed in the cranial sensory ganglia.** *phox2bb* is expressed in the facial, glossopharyngeal, and vagal ganglia (above), and *p2rx3b* is broadly expressed in the vagal ganglia (below). Top, a schematic of an adult fish indicating the regions shown for *phox2bb* (pink polygon) and *p2rx3b* (green rectangle). Both *in situ* images are from adult animals and shown as inverted black and white orthogonal projections. Scale bars in black with scale (microns) indicated. **C. p2rx3b:gfp labels cranial sensory nerves and dorsal root ganglion.** Dorsal root ganglion (DRG) is labeled above, dashed orange inset is magnified center. Organs are indicated above in blue for orientation. Scale bars in white with scale (microns) indicated. Trigeminal (X), facial (VII), glossopharyngeal (IX) and vagal (X) ganglia are indicated in center. Ganglion labeling based on ^30^. The skin generates some speckled autofluorescence, which is marked with a yellow asterisk in center. Purple inset magnified below to show neurite tracing of the vagal ganglia (color code same as panel **E**). Representative image of a 6 month post-fertilization female stained against p2rx3b:gfp with anti-GFP. Across all experiments, we examined p2rx3b:gfp in ∼60 adult animals (both males and females) and found no inter-individual variation in somatotopic arrangement of the vagal ganglia. **D. p2rx3b:gfp expression overlaps with the panneuronal marker HuC/HuD.** Orthogonal projection of immunostained adult expression of p2rx3b:gfp (anti-GFP) co-stained with HuC/HuD (ELAVL). Whole-head image shown at left, white inset is magnified to the right. HuC/HuD alone at top, p2rx3b:gfp in center, channel merge at bottom. The skin generates some speckled autofluorescence, which is marked with white asterisks. **E. Vagal ganglia show stereotyped projection patterns.** Schematic generated by tracing image from Figure 1C, showing innervation targets of the sensory vagus by sub-ganglion.

We report here the first molecular and anatomical characterization of the sensory vagus in adult *D. cerebrum*. We find that the adult *D. cerebrum* sensory vagus contains ∼300 somatotopically organized neurons. Sensory subtypes are highly conserved with those in mouse and express receptors for nutrients, mechanical stimuli, pain, and temperature, along with inflammatory response. We discovered that neurogenesis generates sexually dimorphic left/right asymmetry of the sensory vagus and continues through adulthood. This work lays the foundation for future functional studies on brain-body communication in a small, transparent vertebrate with complex behaviors.

## RESULTS

### Identification of the sensory vagus in adult *D. cerebrum*

To understand the senses that *D. cerebrum* receives from inside of its body, we first sought to identify markers and develop transgenes for the sensory vagus. We used previous studies in zebrafish as a foundation, because the *Danionella* genus split from its sister group (including *D. rerio*) approximately 36 million years ago^28^, allowing the identification of conserved marker genes and the recreation of *D. cerebrum* transgenic lines previously established in zebrafish^21^. Although the sensory vagus in adult zebrafish has not been molecularly studied, larval cranial sensory ganglia express *phox2bb/*DNTS_006240 and *p2rx3b/*DNTS_001674^29,30^. Using hybridization chain reaction (HCR) RNA *in situ*, we found widespread expression of *phox2bb* and *p2rx3b* in adult *D. cerebrum* (**Figure 1B**). Based on a characterized zebrafish transgenic for the cranial ganglia^30^, we established a stable *p2rx3b:gfp* transgenic line in *D. cerebrum*. As in larval zebrafish, we observed expression in the dorsal root ganglion (DRG) neurons in the spinal cord, the trigeminal/facial ganglia (V/VII), the glossopharyngeal ganglia (IX), and the vagal ganglia (X; **Figure 1C**). By co-staining with HuC/HuD(ELAVL), we confirmed that the p2rx3b:gfp transgene labeled most or all of the neuronal cell bodies in the sensory vagus (**Figure 1D**).

The larval zebrafish sensory and motor vagus are arranged in an anterior-posterior layout, such that the anterior-most cell bodies project to the anterior gill arches, and the posterior-most cell bodies project to the posterior gill arches and visceral organs^16,31^. Our *D. cerebrum p2rx3b:gfp* line revealed that this spatial layout is preserved in the adult vagus, with the two anterior-most groups of sensory vagus cell bodies projecting to the gill arches (**Figure 1C,E**). The posterior-most group of neurons, which innervates the seventh gill arch, heart, and viscera^31^ in both zebrafish and *D. cerebrum,* is subdivided into two discrete anatomical clusters of cells: the more anterior and dorsal group projects to the last branchial arch (containing the pharyngeal teeth)^15^, and the most posterior and ventral group projects to the heart and other visceral organs (**Figure 1C,E**). We also observed interspersed neurons projecting to the esophagus, which runs between the gill arches until it reaches the gut. Thus, the *D. cerebrum* sensory vagus can be identified in adult animals by conserved anatomical and genetic features.

### Transcriptional profiling of the *D. cerebrum* cranial sensory neurons

To determine the sensory subtypes represented within the *p2rx3b+* sensory vagus neurons, we used the FLASH-seq full-length single cell RNA sequencing method^32^. We dissected the region containing the sensory vagus from adult male and female animals (48 animals in total), dissociated the cells, and used FACS to isolate *p2rx3b-*GFP+ neurons. 803 cells remained after filtering out low-quality cells (low numbers of features or RNA counts, and excessively high numbers of features, suggesting doublets). This number of cells corresponds to 2-3x coverage, since the sensory vagus contains ∼300 neurons, as based on *p2rx3b:gfp* and HuC/HuD(ELAVL) expression (further detail below). After statistical correction for technical variability^33^ and clustering, we identified 14 cell types (labeled cranial ganglion (CG)0-13).

Cells in all 14 clusters express *p2rx3b* and the canonical sensory vagus marker *phox2bb*, although with differing proportions across the 14 clusters (**Figure 2B**). While studies in mouse and rat have used the expression of the *Phox2b* and *Prdm12* transcription factors to distinguish nodose from jugular vagus, respectively^34,35^, we found that in *D. cerebrum* these two transcription factors are frequently co-expressed in the same clusters (**Figure 2B**). We also found broad expression of the zebrafish cranial sensory nerve markers *entpd3/*DNTS_018855 and *p2rx2*/DNTS_035292^36^ (**Figure 2B**). Conversely, we did not detect expression of the trigeminal and spinal cord marker *p2rx3a*/DNTS_004481^30^, suggesting that we successfully dissected the glossopharyngeal and vagal ganglia away from the broader group of *p2rx3b*-expressing cells (**Figure 2B**).

**Figure 2:**
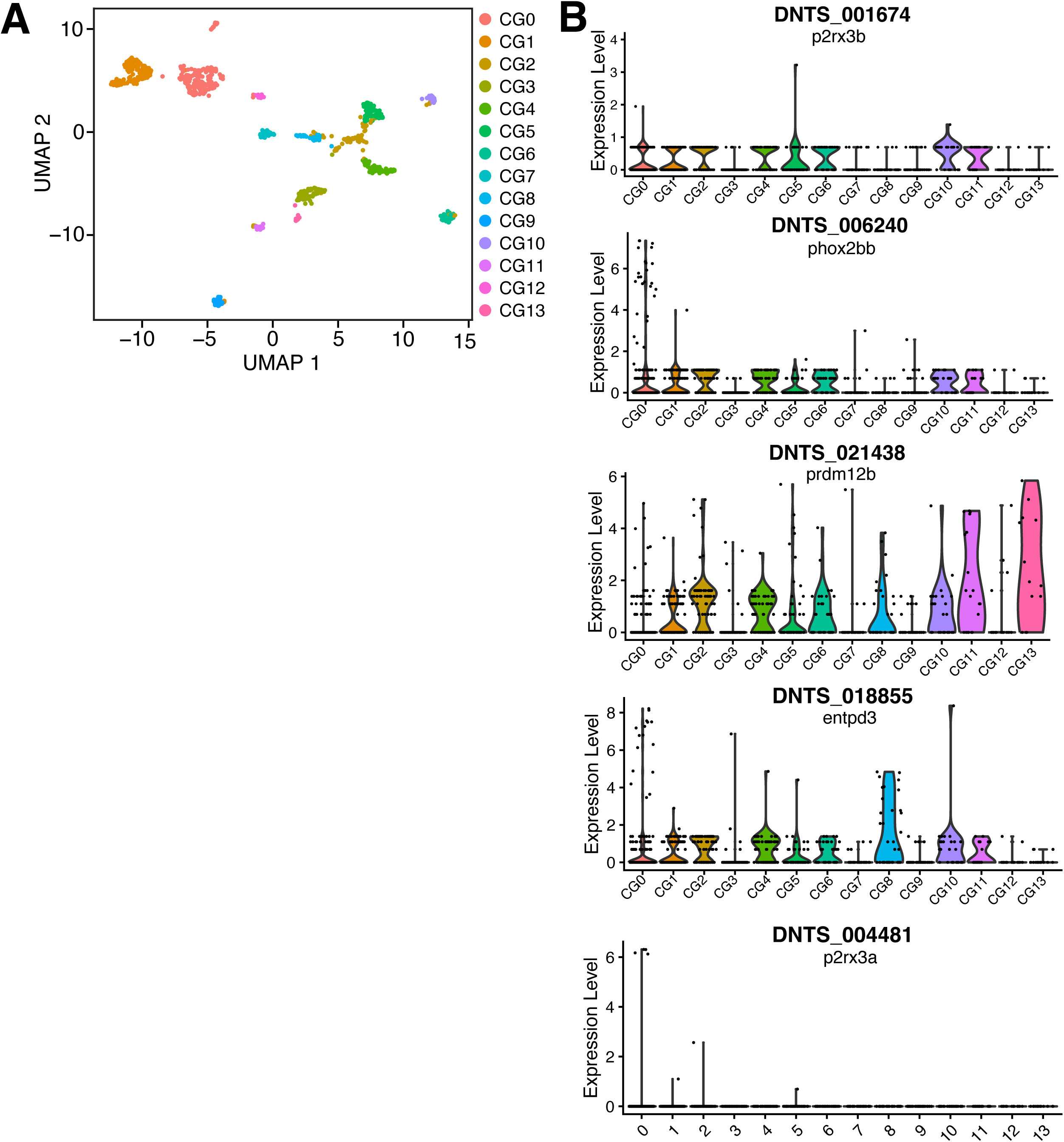
Sequenced p2rx3b:gfp+ neurons can be recognized by broadly-expressed features. **A. p2rx3b:gfp+ cells sort into 14 clusters.** UMAP representation of the sequenced p2rx3b:gfp+ cells. Cells are color-coded by transcriptional cluster **B. Sequenced clusters are broadly marked by known cranial sensory ganglion genes.** Violin plots showing SCT-normalized expression of key cranial sensory marker genes (with both *D. cerebrum* gene annotation name and best zebrafish ortholog) across all 14 clusters. *p2rx3a*, which also marks *p2rx3b*+ trigeminal and spinal cord sensory neurons^30^, is very sparsely expressed.

### Conservation of sensory vagus subtypes

Prior studies in mouse have generated atlases of molecular subtypes in the nodose vagus^1,6,34^. In each case, vagus subtypes and even organ target could be disambiguated using expression of individual or small groups of genes. Thus, we first sought to determine whether we could further distinguish our 14 clusters using established mouse marker genes.

Combining markers from three studies^1,6,34^, we found that all 14 *D. cerebrum* clusters express at least one, and up to 12, key marker genes from the mouse vagus (**Figure 3A**). 12 of the 14 clusters could be uniquely disambiguated using mouse vagal markers alone (**Figure 3B**). Because of the additional whole genome duplication in the teleost lineage in addition to gene-specific duplications^37–40^, many relevant marker genes contain multiple paralogs in fish. For instance, three orthologs for the bioactive lipid receptor Lpar3^6,34^ all have different expression patterns (**Figure 3A**). Other canonical marker genes, such as the mechanosensory Piezo1 (*piezo1*/DNTS_026761) and Piezo2 (*piezo2a/*DNTS_004436 and *piezo2b*/DNTS_032685) are more broadly expressed, limiting their usefulness for resolving individual cluster identities (**Supplemental Figure 1**). We also found that the sensory vagus broadly expresses the Vglut2 paralaog *slc17a6b/*DNTS_011034, but not *slc17a6a/*DNTS_011879 (**Supplemental Figure 2**). These results reveal broad conservation of marker gene expression in the *D. cerebrum* and mouse sensory vagus.

**Figure 3:**
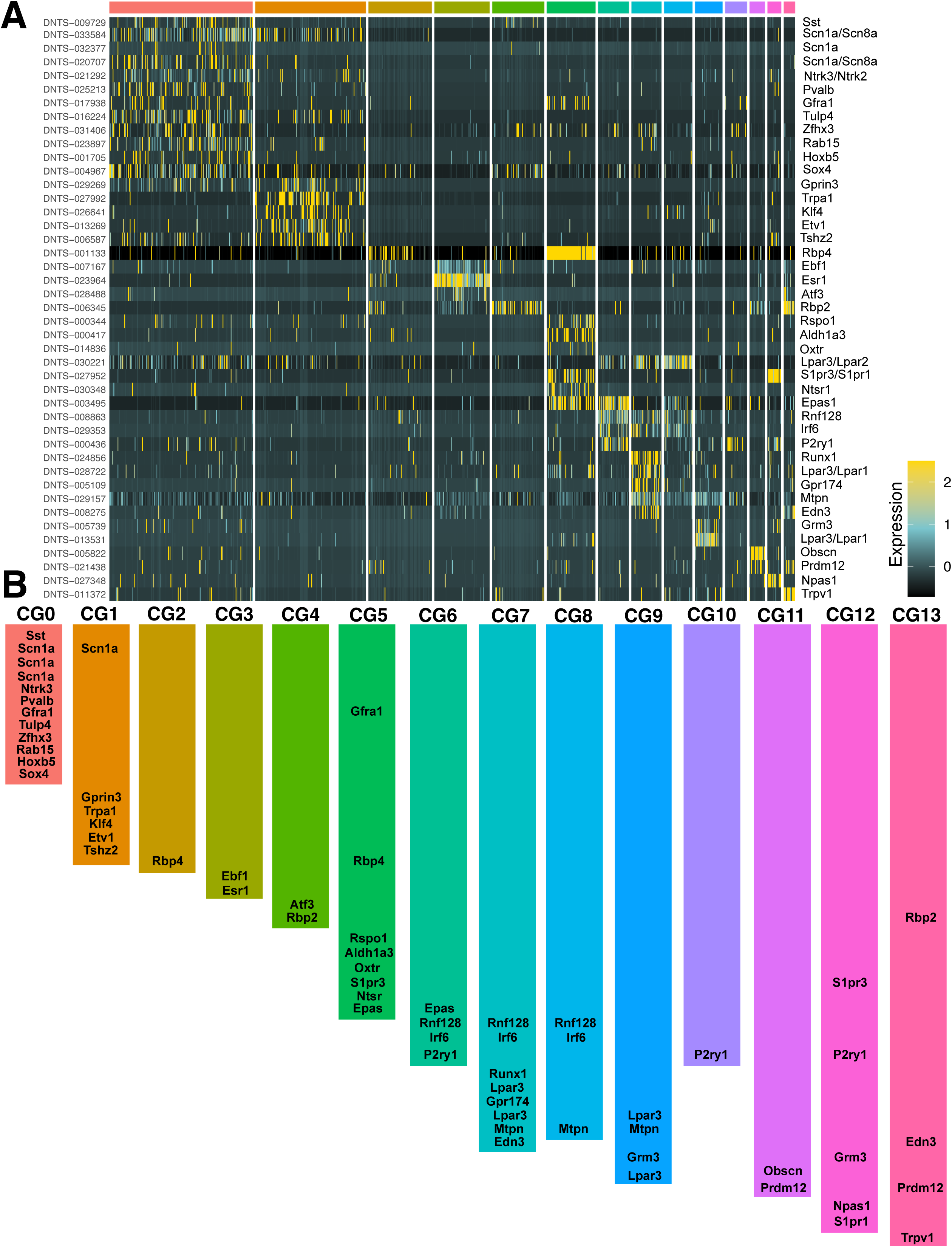
*D. cerebrum* clusters can be disambiguated using mouse marker genes. **A. Heatmap showing expression of mouse marker genes.** Each row shows the expression of one gene, with the *D. cerebrum* gene name shown on the left and the corresponding mouse ortholog shown on the right. After allowing one-to-many orthology, the most diagnostic *D. cerebrum* marker gene was mapped back to the mouse genome, which sometimes returned a better matched one-to-one ortholog than the original marker gene. In this case, both mouse orthologs are shown. Each column corresponds to one cell, with cells broken into the 14 cluster identities (color key at top). **B. 12/14 cranial ganglion clusters are uniquely labeled by mouse marker genes.** Schematic showing which mouse sensory vagus genes are markers of each of the 14 cranial ganglion clusters. All clusters except CG2 (only Rbp4) and CG10 (only P2ry1) are uniquely marked by a combination of mouse marker genes.

### Identification of sensory subtypes

For some clusters we were able to begin assigning sensory modality from conserved mouse marker genes alone, such as the chemosensory cluster CG1 (*Trpa1*) or nociceptive cluster CG13 (*Trpv1*). For other clusters, the conserved markers did not confer sensory modality per se (such as transcription factors), and cluster relationships were not all 1:1. To comprehensively disambiguate and assign sensory modality to our 14 clusters, we extended our analysis from examining conservation with mouse to examining the expression of *D. cerebrum* transcriptional regulators, G-protein coupled receptors (GPCRs), and transient receptor potential (TRP) channels. We found that transcriptional regulators and GPCRs were each individually sufficient to identify all clusters uniquely (**Figure 4A,B**). Although much smaller, the TRP channel family also provided unique marker genes for most clusters (**Figure 4C**). We were able to identify a variety of putative sensory modalities (mechanosensory, chemosensory, thermosensory, nutrient-sensing, nociceptive, and inflammatory response), and combinations of polymodal classes (**Figure 4D**).

**Figure 4:**
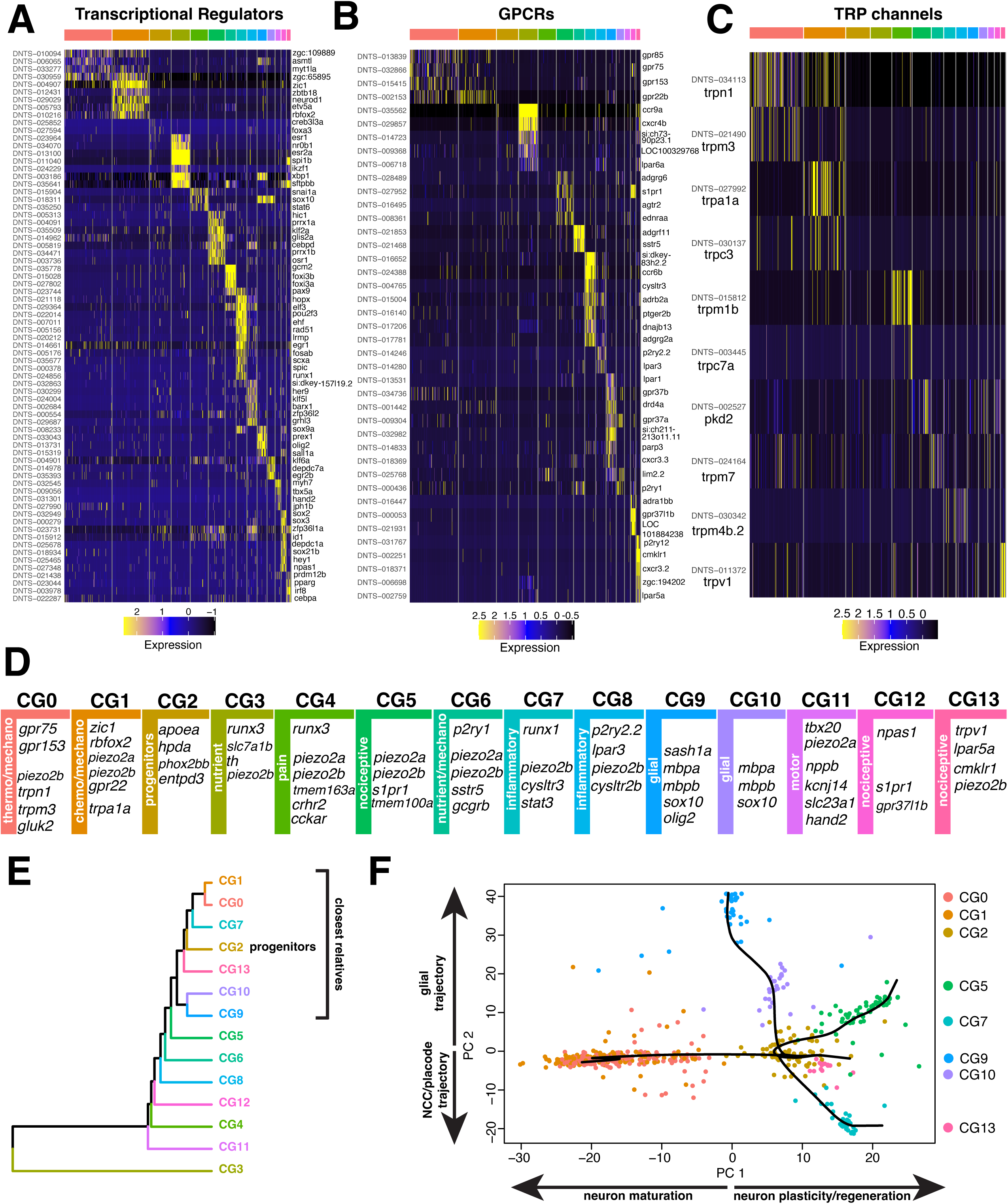
Transcriptional clusters can be assigned functional identities based on key marker genes. **Transcriptional regulators (A), GPCRs (B), and TRP channels (C)** are differentially expressed across the 14 cranial ganglion clusters. Genes of each class were extracted from the zebrafish genome using BioMart, matched with the top *D. cerebrum* ortholog, and then filtered against statistically significant cluster markers. **D. Cellular identities of transcriptional clusters can be assigned based on marker genes.** Proposed identities of each of the 14 transcriptional clusters are shown, along with key significantly enriched marker genes that were considered diagnostic for assigning cellular identity. **E. The CG2 progenitor cluster has several phylogenetic relatives.** Unrooted cluster tree of phylogenetic relatedness among the 14 transcriptional clusters. Closest relatives to CG2 were selected based both on the shared node in the phylogenetic tree, and a cophenetic distance cutoff (25,000). **F. CG2 has potential trajectories to multiple other transcriptional clusters.** Pseudotime relationships are shown between CG2 and its closest relative clusters. Trajectories are plotted in PC space. The top marker genes for each PC were used to assign differentiation trajectories for each lineage, shown on the arrows along each axis.

#### Polymodal (CG0, CG1, CG6)

The presence of polymodal sensory neurons has emerged as a recurring theme in studies of the mouse sensory vagus^1,2,34^. We determined that three clusters (CG0, CG1, and CG6) contain neurons expressing receptors capable of responding to both mechanical cues and an additional stimulus, which was different for each cluster (thermosensitive, chemosensitive, or nutrient-sensing).

CG0 expresses both the mechanosensory *trpn1/*DNTS_034113 and noxious heat-sensing^41^ *trpm3*/DNTS_021490, although in mostly non-overlapping individual cells. Interestingly, *gpr75/*DNTS_032866 is also expressed, which is known to function in the nervous system to control appetite-regulation and energy balance, and has thermogenic phenotypes^42^. Unexpectedly, our dataset also gave insights into cold-sensing receptors in teleosts. Cold-adapting and -responding neurons were recently described in larval zebrafish^43^. However, as the canonical TrpM8 cold receptors have been evolutionarily lost from teleosts^44^, the molecular receptor such neurons may be utilizing is unknown. We found that CG0 also expresses the glutamate receptor *gluk2/*DNTS_028951. This glutamate receptor is broadly conserved across animals and has been demonstrated in *C. elegans* and mouse to serve as a cold receptor, with a channel temperature threshold of ∼18C ^45^. In zebrafish with *gluk2* mutations, locomotory response to a uniform 18C temperature is diminished^46^. Taken together this suggests that CG0 contains mechanosensory and thermosensory neurons competent of sensing both cold and hot temperatures, and may intersect with appetite regulation.

CG1 is marked by *gpr22*/DNTS_002153, which has been found in gut-innervating nodose vagus neurons in mouse^47^. Expression of the chemosensory *trpa1a* and of the transcription factors *zic1/*DNTS_004907 and *rbfox2*/DNTS_010216 distinguish CG1 from CG0. Rbfox2 is a known marker of cranial and dorsal root ganglia in mouse^48^. Thus, CG1 contains mechanosensory and chemosensory neurons.

CG6 expresses the *p2ry1* purinergic receptor. *p2ry1+* mechanoreceptive vagal neurons are well-established in mouse^5,34^, and indeed we found that CG6 expresses *piezo2a* and *piezo2b*, suggesting a mechanosensory role. CG6 is also distinguished by strong expression of the *sstr5/*DNTS_021468 somatostatin receptor, and expression of the nutrient-sensing *gcgrb*/DNTS_006318 glucagon receptor. These results suggest that CG6 is both mechanosensory and nutrient-sensing.

#### Nociceptive (CG5, CG12, CG13)

An additional three clusters (CG5, CG12, and CG13) were identified by their expression of known nociceptive receptors and channels.

CG5 and CG12 both express the nociceptive receptor^49^ *s1pr1*/DNTS_027952. CG5 also expresses the transmembrane protein *tmem100a*/DNTS_022136, which is required for inducing mechanical sensitivity in nociceptors in response to inflammation^50^. CG12 is distinguished by strong expression of the epibranchial placodal marker^51^ *sox3/*DNTS_000279, and *npas1/*DNTS_027348, which has been identified as a marker for subdiaphragmatic nodose vagus in mouse^6^. CG13 is unique in its strong expression of both *trpv1*/DNTS_ 011372 and *lpar5a*/DNTS_002759, suggesting that it also contains nociceptive neurons. It also expresses *cmklr1*/DNTS_002251, a receptor for the chemerin signaling axis, which has been implicated in baroregulation^52^.

#### Inflammatory response (CG7, CG8)

Sensory vagus neurons have direct access to potentially harmful stimuli (e.g. via ingestion or inhalation) and thus are poised to initiate systemic inflammatory responses. Two transcriptional subtypes, CG7 and CG8, were best characterized by their expression of inflammatory receptors.

In addition to the *runx1/*DNTS_024856 somatosensory marker, CG7 expresses *cysltr3*/DNTS_004765, which has been implicated in avoidance behaviors caused by re-exposure to gastrointestinal allergens^53^. CG7 also expresses *stat3*/DNTS_035246, which is expressed in neurons mediating anti-inflammatory responses in both the motor vagus and DRGs^54,55^, and in cranial ganglia in zebrafish^56^. CG8 expresses a different subset of inflammatory response genes, *cysltr2*/DNTS_ 021959 and *lpar3*/DNTS_014280.

#### Nutrient-sensing (CG3)

CG3 expresses the *runx3* somatosensory marker and is strongly marked by tyrosine hydroxylase *th/*DNTS_024425, but not *dbh*, consistent with vagal dopaminergic neurons that have been identified in other vertebrates^57^. CG3 also expresses *slc7a1b*/DNTS_023138. *slc7a1b*, also known as CAT1 in vertebrates and mosquitoes, and *slif* in Drosophila, has been shown to directly transport amino acids, and functions as a nutrient sensing mechanism in Drosophila and *A. aegypti*^58,59^. Combined, CG3 may represent nutrient sensing neurons.

#### Pain (CG4)

CG4 expresses *tmem163a/*DNTS_016371, which is known to modulate purinergic (P2X) signaling in pain-sensing neurons^60^, and *crhr2*/DNTS_014811, which is involved in stress responsive and also marks the mouse nodose vagus^47^. CG4 also expresses the *runx3*/DNTS_025958 transcription factor, which is a conserved marker of somatosensory neurons between mammals and zebrafish^61–63^.

#### Motor (CG11)

CG11 appears to be a cluster of cranial motor, rather than cranial sensory neurons, based on the strong expression of *tbx20*/DNTS_015167 and *kcnj14*(Kir2.4)/DNTS_030377^64,65^. In larval zebrafish, transgenic lines that predominantly label vagal motor neurons (which have cell bodies in the hindbrain rather than the peripheral ganglia) also label a subset of neurons in the peripheral vagal ganglion^16,31^. Because the peripheral vagal ganglion was assumed to be entirely composed of sensory neurons, it was thought that these transgenic lines may label both cranial motor neurons and a subset of peripheral sensory neurons. However, there might indeed be a population of vagal motor neurons that coexists in the peripheral sensory ganglion.

#### Neural progenitors (CG2)

CG2 was initially the least decipherable cluster. Unlike every other cluster, it does not have a unique signature of either transcriptional regulators (**Figure 4A**) or GPCRs (**Figure 4B**). A recent whole-body sequencing study of *D. cerebrum* identified a transcriptional subpopulation that resembles Schwann cell precursors (SCPs) and multipotent neural-crest-like cells^66^. A single subcluster of this population was shown to express *phox2bb* and *entpd3*, consistent with zebrafish cranial sensory nerves and our CG2 specifically. We were also able to corroborate the expression of other markers of this reported SCP progenitor cluster, such as *apoea*/DNTS_029317 and *hpda*/DNTS_003290, and the pre-placodal marker *six1a*/DNTS_001639^67^. Thus, we interpret CG2 as an adult progenitor population contributing to the cranial sensory nerves.

We were curious whether any other clusters represented transcriptional descendants of CG2. We performed phylogenetic analysis to identify the transcriptionally most similar clusters to CG2, and then used pseudotime analysis to identify potential trajectory relationships between CG2 and nearby clusters(**Figure 4E,F**). We found that these proposed pseudotime trajectories were well-represented by the relationship of the clusters in principal component space, where CG2 is located near the center and the related clusters extend out along PCs 1 and 2. The trajectory between CG2 and CG0/CG1 follows a principal component defined by neuronal maturation genes (*snap25a*/ DNTS-014883, *sncb*/DNTS-031200, *stxbp1a*/DNTS-025903). A trajectory between CG2 and CG5 follows this principal component in the opposite direction, defined by *krt4*/DNTS_ 026585 (a marker of neural crest-derived regeneration in zebrafish^68^) and *eef1a2*/DNTS_032506 (required for neuron plasticity and survival^69,70^). A trajectory also exists between CG2 and CG7 along the second principal component, defined by the placodal differentiation markers *epcam/*DNTS_ 029664 and *elf3*/DNTS_029364^67^, and the neural crest differentiation marker *krt18a.1*/DNTS_027834^71^. In the opposite direction, the CG9 and CG10 PC is defined by canonical Schwann cell glial markers such as *mbpa*/DNTS_022628, *mbpb*/DNTS_006991, and *plp1b*/DNTS_000421 ^72^. The presence of differentiation trajectories leading to both new neurons and to new glial cells is consistent with an SCP identity for CG2.

### Neurogenesis in the adult sensory vagus

Teleosts generally have a high degree of adult neurogenesis^73^, but continuous neurogenesis in peripheral neurons has also been identified in other vertebrates^74^. In the periphery, new neurons can be generated either via stem cell-derived neural progenitors^75^, or via divisions of *sox10+* Schwann precursor cells (SCPs)^76,77^. As *sox10* was a key marker for our CG2 progenitor cluster, we examined its localization in adult *D. cerebrum*. At 9 months post-fertilization, SOX10+ cells are abundant within the vagal ganglion, and arrayed alongside the processes (**Figure 5A**), suggesting that SOX10+ cells persist throughout adulthood as a potential source of new vagal neurons.

**Figure 5:**
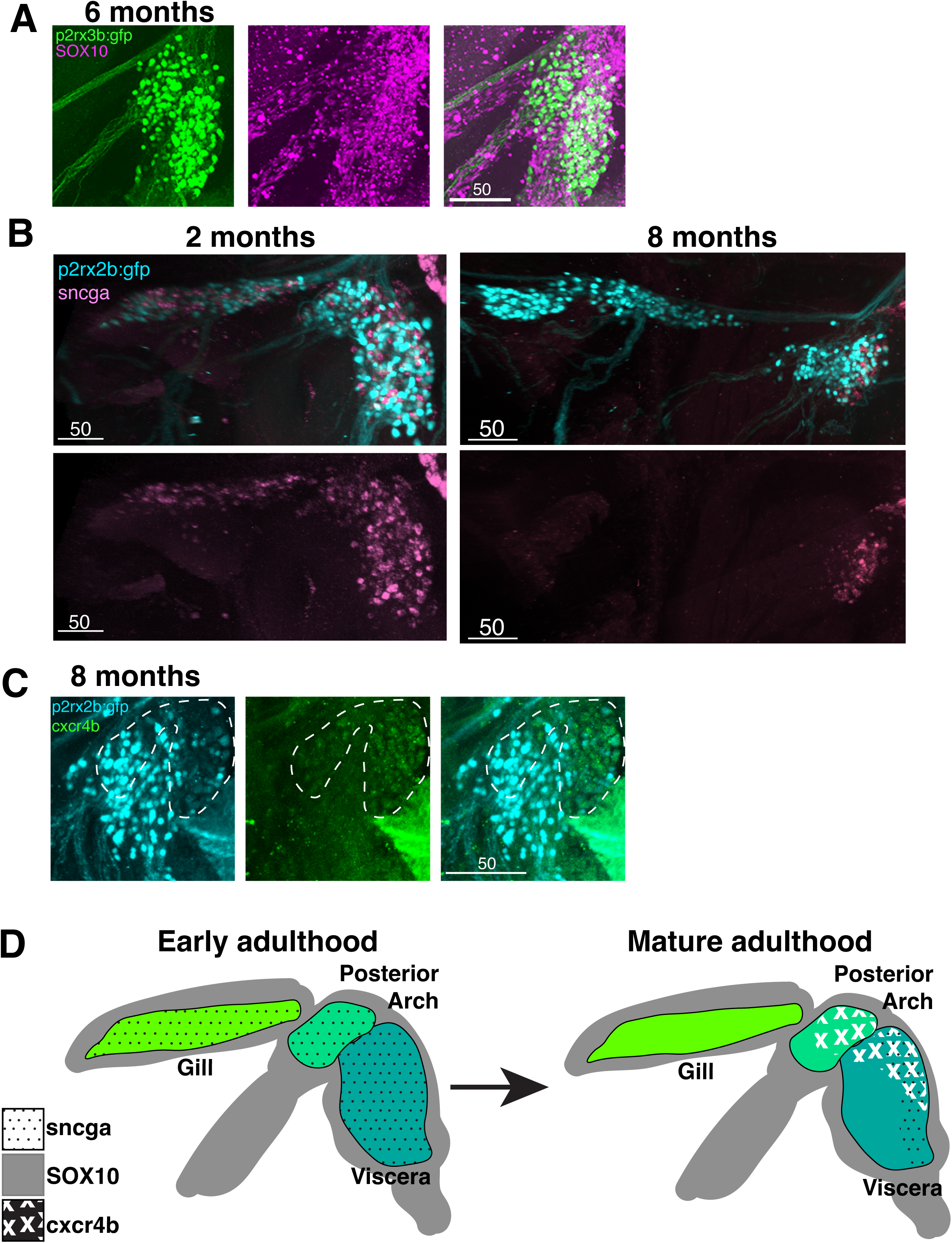
Widespread adult neurogenesis becomes spatially restricted with age. **A. SOX10+ cells are present within and around the vagal ganglion.** Anti-GFP (p2rx3b:gfp) and anti-SOX10 staining of an adult (6 months post-fertilization) animal. Orthogonal projections of p2rx3b:gfp in green on left, SOX10 in magenta in center, and merged stack on right. SOX10+ cells are especially found on the periphery of the ganglion and along the axon tracts. Scale bars in white with scale (microns) indicated in all panels. **B. *sncga+* cells are initially widespread and then become restricted.** *sncga* transcription (HCR RNA *in situ*) at 2 months (left) and 8 months (right) post fertilization, with *p2rx3b:gfp* (anti-GFP staining) for anatomical reference. Above, merged orthogonal projection of *sncga* and *p2rx3b:gfp,* below, *sncga* single channel image. **C. *cxcr4b+* cells are present at the dorsal and posterior edge of the vagal ganglion.** HCR RNA *in situ* of *cxcr4b* with *p2rx3b:gfp* (anti-GFP staining) for anatomical reference at 8 months post fertilization. Orthogonal projection of *p2rx3b:gfp* alone left, *cxcr4b* alone center, and merged channels image right. White dashed outline indicates the proliferative dorsal and posterior edges of the visceral ganglion. **D. Schematic summary of sites of vagal neurogenesis across adulthood.** “Early adulthood” designation is based on our data from 2 months post fertilization, “Mature adulthood” is based on our data from 6-8 months post fertilization.

We next turned to genes that are associated with neurogenesis of the cranial sensory ganglia in larval zebrafish. The synaptic scaffolding gene synuclein-gamma/*sncga* is expressed in the larval zebrafish cranial sensory ganglia and has been implicated in placodal sensory axon outgrowth^78,79^, and was a marker of our progenitor subtype. We found that in earlier adulthood (2-4 months post fertilization), *sncga* is very broadly expressed across the sensory vagus (**Figure 5B**), reminiscent of larval zebrafish. Most of these cells are *p2rx3b:gfp-,* although some show weak *p2rx3b:gfp* expression, suggesting that they may be nascent neurons. In 8-month old adults, the expression of *sncga* has become restricted to the most posterior and ventral region of the vagal ganglion, in the location of neurons that project to the visceral organs (**Figure 5B**). It is still expressed in both *p2rx3b:gfp-* and weakly *p2rx3b:gfp+* cells, suggesting that neurons are still being generated.

We also examined expression of the GPCR *cxcr4b*/DNTS_029857, which in zebrafish is expressed in cranial sensory placodes^80,81^. In the olfactory placode, *cxcr4b* is expressed broadly during early neurogenesis, and then becomes restricted to one edge of the ganglion^80^. Expression of *cxcr4b* in the sensory vagus in 8-month old adult *D. cerebrum* was reminiscent of this peripheral zebrafish expression: *cxcr4b+* cells were numerous on the posterior and dorsal edges of the ganglion, but sparse or absent in the rest of the ganglion (**Figure 5C**). This pattern of *cxcr4b+* cells was also consistent with the progressive restriction of *sncga* expression in the sensory vagus (**Figure 5D**). As SOX10 remains widespread, it is possible that there are additional neurogenic lineages, but we conclude that the *cxcr4b+* and *sncga+* trajectories in the vagal ganglion become spatially restricted as adulthood progresses.

### Anatomical organization of molecular subtypes

We took advantage of the somatotopic zonation of the ganglion to spatially map transcriptional sensory subtypes. Some neuron types were distributed across the entire vagal ganglion, in both the gill- and viscera-innervating regions. These included neurons expressing the neuropeptide *calca/*DNTS_014345, which marks stomach-projecting vagal neurons in mouse^6,34^, and the mechanosensory *piezo2b/*DNTS_032685 (**Figure 6B**). Other neuron types were restricted to particular spatial regions. The *gluk2* glutamate receptor was only expressed in the cluster of vagus neurons that project to the most posterior gill arch (**Figure 6D**). Multiple genes are also specific to the viscera-innervating neurons, such as the *trpa1a*/DNTS_027992 chemosensory channel and the mechanosensory *piezo2a/*DNTS_004436/DNTS_004431 (**Figure 6C**). This *trpa1a* expression pattern matches larval zebrafish, where the *trpa1b* paralog is more broadly expressed in the cranial sensory ganglia^82^. We found that this difference in breadth was also the case for the *piezo2* paralogs, where in contrast to the viscera-specific *piezo2a, piezo2b* is expressed across the entire vagus sensory ganglion (**Figure 6B,C**).

**Figure 6:**
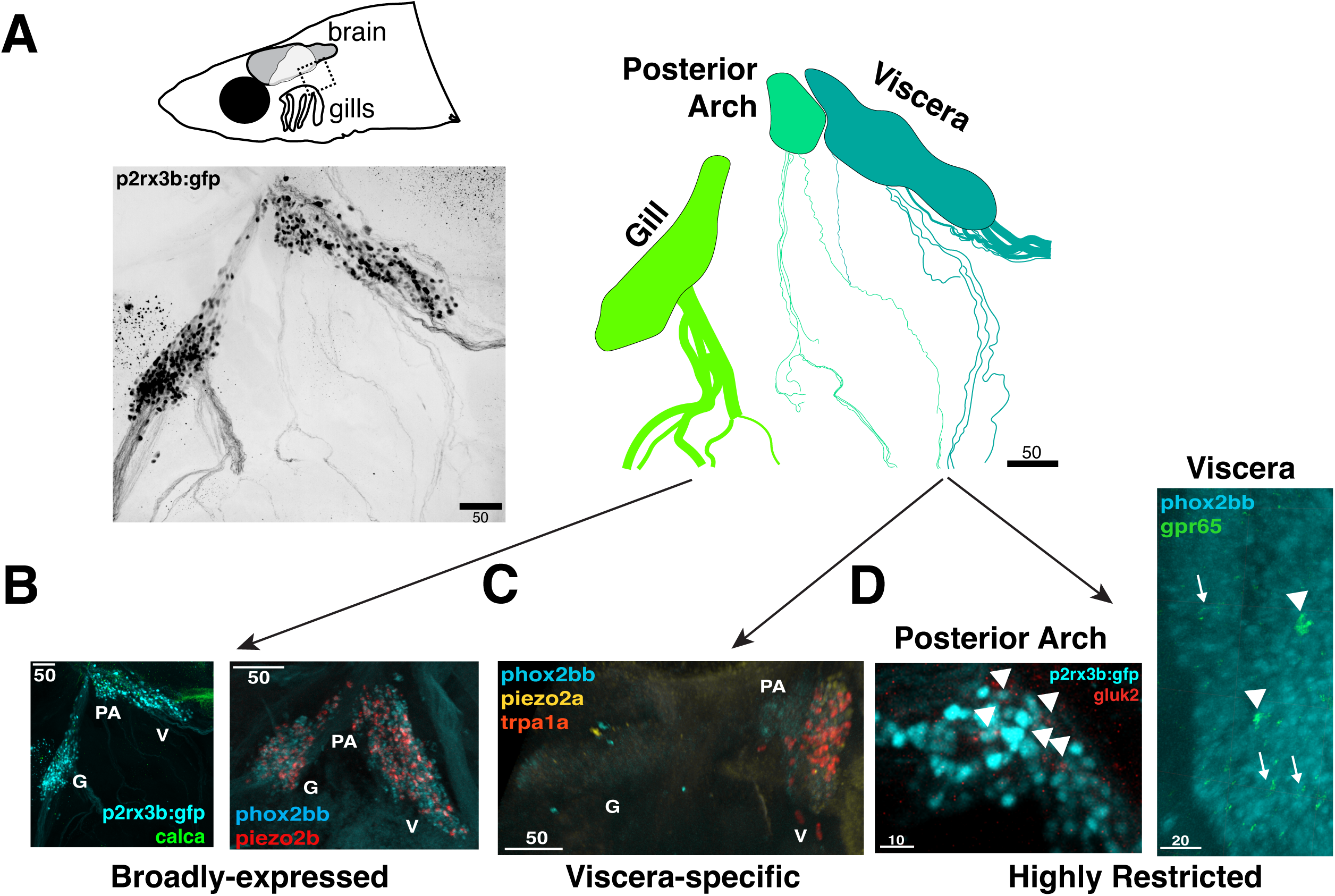
Molecular subtypes are spatially restricted in the vagal ganglion. **A. The vagal ganglion can be subdivided based on innervation target.** p2rx3b:gfp-expressing neurons (left) have stereotyped innervation targets based on where they reside in the ganglion (schematized tracing of neuronal projections, right). Black and white inverted image of anti-GFP-stained p2rx3b:gfp in an adult animal. Scale bar in black with scale (microns). Schematic fish above indicates anatomical location of image (black dashed box). **B. Broad subtypes show expression in the gills, posterior arch, and viscera-innervating regions.** Representative images of HCR RNA *in situ* for the *calca* neuropeptide with p2rx3b:gfp as anatomical reference, and *piezo2b* mechanosensory channel with *phox2bb* as anatomical reference. In all images: G, gill; PA, posterior arch; V, viscera. Scale bars in white with scale shown (microns) in all images. (The *calca* image is the same animal from Figure 6A, as this animal was stained for both the p2rx3b:gfp transgene and HCR in situ). **C. Some subtypes are restricted to the viscera-innervating ganglion.** HCR *in situ* of the *piezo2a* mechanosensory channel and *trpa1a* chemosensory channel, with *phox2bb* as a reference. **D. Some sensory receptors are only expressed in a few vagal neurons.** HCR *in situ* of the gluk2 glutamate receptor with p2rx3b:gfp as an anatomical reference, and the *gpr65* nutrient receptor with *phox2bb* as an anatomical reference. Images are zoomed to the relevant regions (posterior arch ganglion for *gluk2* and visceral ganglion for *gpr65*) to show expression. *gluk2+* cells are indicated by arrowheads. The arrowheads in the *gpr65* image indicate high *gpr65* cells, and the arrows indicate examples of low *gpr65* cells.

The acid-sensing *gpr65/*DNTS_006003 also showed regional differences in expression level. Many cells across the ganglion have low expression (1-2 punctae per neuron), but 3-4 neurons show high expression and are restricted to the cluster of vagus neurons that project to the viscera (**Figure 6D**). This is reminiscent of studies in the mouse sensory vagus, where high- and low-expressing populations of Gpr65+ neurons (which mark subpopulations of nutrient-sensing neurons^2^) have also been identified^34^. Consistent with studies in mouse^34^, we find that these high *gpr65* neurons also express *piezo2b,* suggesting that they are polymodal (**Supplemental Figure 3**). Taken together, we find that the composition of sensory subtypes differs by anatomical region, suggesting that the gills and internal organs are capable of responding to different sensory information.

### Sexual dimorphism in the adult sensory vagus

*D. cerebrum* has the smallest brain of any known vertebrate, with ∼650,000 neurons^21^ compared with ∼10 million in an adult zebrafish^22^. In mouse, the number of neurons in the sensory vagus has been estimated at 2500 per ganglion (5000 in total)^83^. We used our p2rx3b:gfp transgene to count the number of cell bodies in the sensory vagus in adult *D. cerebrum*. We counted ∼150 p2rx3b:gfp+ neurons on both the left and right sides (**Figure 7A**). Taking estimated number of neurons in the brain as a denominator, this suggests that *D. cerebrum* has an order of magnitude higher proportion of sensory vagus neurons relative to mouse, 0.05% (300/650,000^21^) vs. 0.007% (5000^83^/70,000,000^84^).

**Figure 7:**
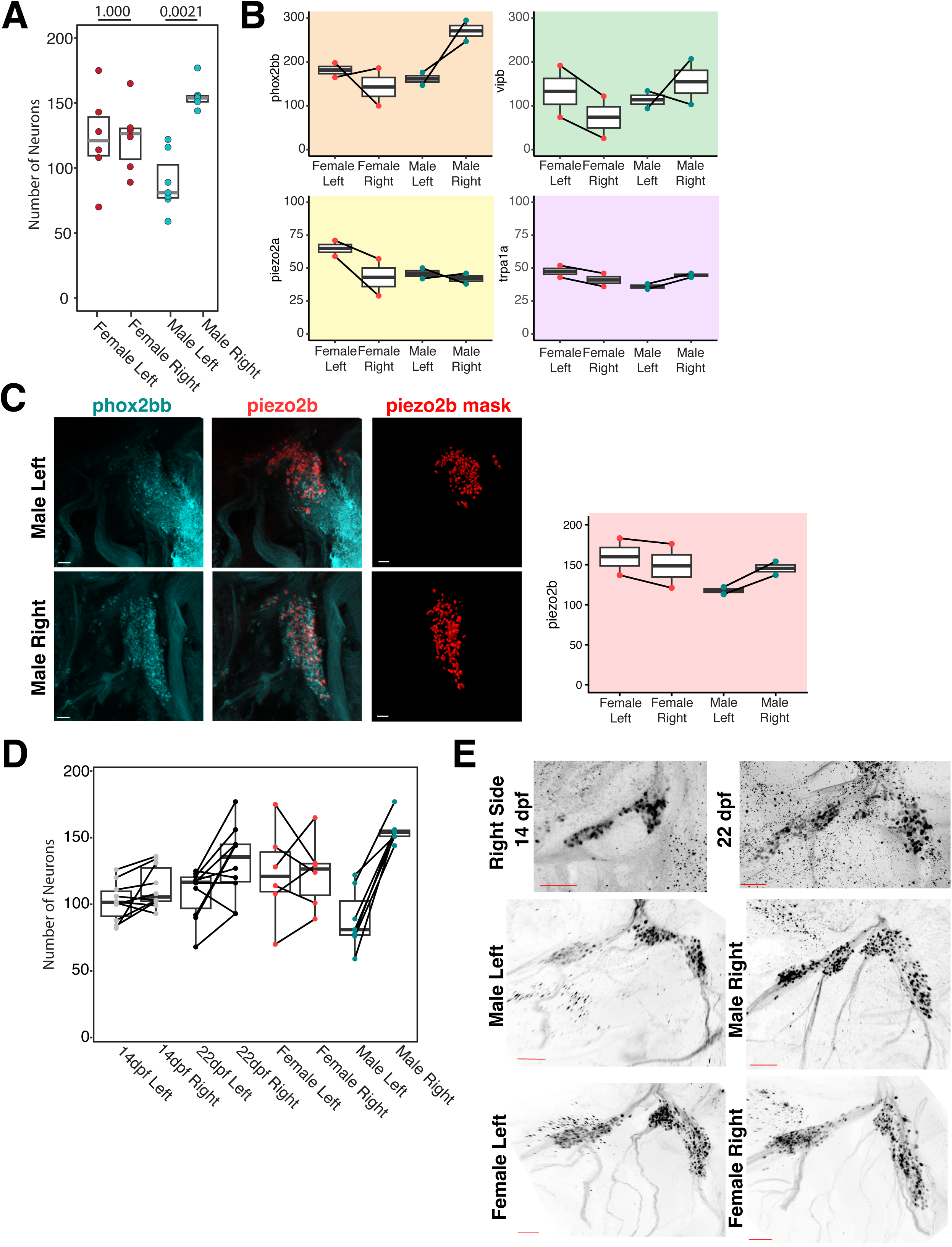
The sensory vagus is asymmetric in adult male animals. **A. *p2rx3b+* neuron numbers are asymmetric in the male vagus.** Beeswarm plot showing number of GFP+ neurons. Each dot represents the number of neurons from one side of individual animal, gray bars show the median, black boxes show quartiles. p-values shown at top of chart calculated by Wilcoxon rank-sum test. **B. There is not a general expansion of sensory subtypes on the male right side.** Quantification of *phox2bb*, *vipb, piezo2a,* and *trpa1a* HCR RNA *in situ* signal in adult males and females on the left and right sides. Although *phox2bb* expression recapitulates the asymmetry of *p2rx3b:gfp*, none of the other subtype markers are asymmetric. RNA *in situ* signal was quantified using automated masking (see Methods), y axis shows number of masks (not directly representative of number of neurons). Each dot represents the count from one animal, solid bar shows median, black box shows quartiles. **C. Males have more piezo2b vagal neurons on the right side than left side.** Left, HCR *in situ* showing *phox2bb, piezo2b,* and masked *piezo2b+* neurons (see Methods) in an adult male animal. Scale bars (white) are 20 microns. Right, quantification of *piezo2b* neuron masks in adult males and females on the left and right sides. **D. The vagus is symmetric in juvenile animals.** Beeswarm plot showing number of GFP+ neurons in 14dpf, 22dpf, and adult fish. The adult male and female neuron counts are the same data from Figure 7A, shown again here in the context of juvenile neuron counts. Each dot represents the number of neurons from one side of individual animal, black bars show the median, black boxes show quartiles. Diagonal black lines connect the left and right side neuron counts from each individual animal. **E. Sensory vagus dimorphism is generated developmentally.** Anti-GFP staining of p2rx3b:gfp in 14dpf, 22dpf, and 2 month old (male and female) animals to demonstrate increase in neuron count and adult male asymmetry. GFP signal is shown in color-inverted grayscale. Scale bars (red) all 50 microns. Female Right image is the same animal from Figure 6A.

Surprisingly, when quantifying p2rx3b:gfp+ neurons, we found that the number differed between adult male and female animals. Specifically, males have significantly more neurons on the right side of the sensory vagus than the left side (median 81 neurons on the left side, and 154 neurons on the right side), whereas females have no asymmetry (median 121 neurons on the left side, and 127 neurons on the right side) (**Figure 7A,D**).

We sought to determine the identify of these additional right-side neurons. We found that none of our molecular subtypes were male-specific. In addition, the male asymmetry is not a general expansion of all sensory subtypes on the male right side: for instance, we found the same number of *trpa1a*, *piezo2a*, and *vipb* neurons between males and females, and the left and right sides (**Figure 7B, Supplemental Figure 4**). In contrast, expression of the light touch receptor *piezo2b* in the visceral vagal ganglion recapitulated the male-specific asymmetry (**Figure 7C**). These observations indicate that there is an increased number of neurons from multiple molecular subtypes on the male right side.

Because of the evidence for adult vagal neurogenesis in our RNA sequencing data, we wondered when in development this asymmetry was established. In zebrafish, the gonad and germ cells (and entire organism) are initially bipotential, and begin to sexually differentiate via a stepwise process of gene expression, cell division, and cellular differentiation between 12dpf and 21dpf^85^. By anatomical criteria alone in *D. cerebrum,* we were similarly unable to distinguish male from female gonads at 14dpf, and thus considered this a timepoint before somatic sexual maturation is apparent. The number of p2rx3b:gfp+ sensory vagus neurons on the left and right sides of 14dpf animals were near-perfect symmetry, with a median number of 102 neurons on the left side, and 103 neurons on the right side (**Figure 7C,D**). Consistent with our observations that general neurogenesis is also occurring, the median number of neurons in 14dpf animals was fewer than the median number we counted in adults (on either side or in either sex). At 22dpf 9 of 10 animals still had ambiguous gonads, and 1 of 10 was discernibly female. Thus, we considered this to be a timepoint of active sexual differentiation. The number of p2rx3b:gfp+ cells mirrored this interpretation: the median number of neurons on the left side was now 117, and on the right side was 136 **(Figure 7C,D**). This asymmetry in the median is driven by a subset of animals, as is expected if it is only developing in one sex. We conclude that male asymmetry begins to be established during the third week post-fertilization, before clear anatomical differentiation of the gonad, by extra right-sided neurogenesis of sex-shared molecular subtypes.

## DISCUSSION

We present here a characterization of transcriptional subtypes, anatomical organization, and sexual dimorphism in the sensory vagus ganglion of adult *D. cerebrum*. One theme that emerged from this analysis is the deep evolutionary conservation between *D. cerebrum,* zebrafish, and mouse. Comparing *D. cerebrum* adults and larval zebrafish, we found high conservation of anatomy, both in regard to the physical arrangement of the ganglia and to the somatotopic projection pattern ^15,16,30,36^. We also found conservation compared to larval zebrafish of genes that are both broadly-expressed (e.g. *p2rx3b*, *phox2bb*, and *entpd3*) or more restricted in the cranial sensory ganglia (e.g. viscera-specific expression of *trpa1a*). While molecular analysis has not been extended to adult zebrafish, our results suggest that this conservation will hold across developmental stages.

Since the number of defined subtypes in the mouse sensory vagus differs between different studies^1,2,5,6,11,12,34,86,87^, we focused on evaluating conservation using marker genes, which are more consistent across studies. We were able to disambiguate 12 of our 14 transcriptional clusters using mouse marker genes alone. These diagnostic marker genes cover a variety of gene families (sensory receptors and channels, transcription factors, cytoskeletal components), suggesting a breadth of shared characteristics.

We found a preponderance of Piezo expression across molecular subtypes (we identified 11/14 *piezo1+* clusters, 8/14 *piezo2a+*, and 10/14 *piezo2b+*), suggesting major mechanosensory roles of vagus neurons in *D. cerebrum*. Analogously, expression and functional analyses have demonstrated that many neurons in the mouse sensory vagus are mechanosensory across multiple target organs^1,2,6,11,12,34,88^. These neurons are capable of responding to physiological stimuli as diverse as gut distension^2,6^, arterial blood pressure^88,89^, or laryngeal/airway stretch^11^. Functional analysis in teleosts has suggested that some of these roles of stretch-sensitive neurons are likely shared. For example, in larval zebrafish it has been demonstrated that neurons in the sensory vagus respond to changes in heart rate^20^. While the sensory stimulus used to detect heart rate has not been described in larval zebrafish, we propose that this is one role of stretch-sensitive neurons in the *D. cerebrum* sensory vagus.

In contrast to the conserved broad expression of mechanosensory receptors, we only identified two of our clusters as potentially nutrient-responsive (CG3 and CG6), whereas in mouse several classes of nutrient-activated mucosal afferents exist^6^. A recent study in larval zebrafish found that while amino acids promote feeding and activate vagal neurons, glucose is a neutral stimulus for zebrafish (in contrast to mouse)^17^. Correspondingly, our nutrient-responsive subtype CG3 expresses the amino acid transporter *slc7a1b*. The smaller complement of nutrient-sensing neurons also reflects the simpler anatomy of the fish gut. For example, the zebrafish gut is a linear tube without precise boundaries between subregions^90^, whereas different gut regions are innervated by distinct groups of neurons in the mouse^2,6^. The combination of anatomical simplification of the teleost gut and reduction in nutrient discrimination may underlie the smaller complement of functional gut-innervating classes compared to mouse.

We also identified neurons with thermosensitive receptors capable of responding to both warm and cool temperatures (CG0). *D. cerebrum* has been found to prefer water depths where the temperature is around 25C^91^, suggesting that temperature is a physiologically relevant stimulus. Mammals have numerous thermosensitive vagal afferents that are thought to respond to both environmental and internal temperature.

For example, the bronchospasm in response to inhalation of cold air is abolished by vagotomy ^92^. Mammals also perform behavioral thermoregulation in response to systemic stimuli such as fever^93^. The *D. cerebrum* thermosensory neurons might thus also respond to internally generated temperature changes.

Several of the remaining sensory clusters we identified express genes related to nociception and inflammatory response (CG5, CG7, CG8, CG12, CG13). Some of these cells might serve respiratory protection. For example, the vagal sensory ganglion in teleosts also innervates the more posterior gill arches^14^. While the mammalian lung and gills are not evolutionarily homologous^94^, both receive sensory and motor innervation from the vagus nerve. Electrical stimulation of the vagus nerve in fish leads to coughing and changes in respiratory rhythm^95^. Detailed characterization of the sensory vagus in the mouse lung has revealed that, despite belonging to several different sensory types, all subtypes express genes responsive to chemical irritants, acids, and/or inflammatory cues^12^. In *D. cerebrum*, the preponderance of nociceptive and inflammatory response genes we identified may relate to an evolutionarily shared goal of respiratory protection.

In summary, we conclude that sensory vagus neurons in *D. cerebrum* and mouse share a variety of molecular features, such as numerous mechanosensory subtypes, inflammatory response neurons, and thermosensitive neurons. They differ, however, in the number of nutrient-sensing subtypes, and in anatomy of target organs (such as gills vs. lungs, or differences in gut compartmentalization).

We also identified anatomical asymmetries of the adult male *D. cerebrum* vagus nerve. We found that males have an increased number of *piezo2b+/p2rx3b+* neurons on the right side of the sensory vagus. Anatomical studies in rats have also identified left/right differences in the projection patterns of both the sensory and motor components of the vagus nerve to the digestive system^96,97^, suggesting that vagal asymmetry affects multiple organ systems. Additional molecular left/right asymmetries have been more recently described in vagal sensory neurons^98^. For instance, it has been shown that there is lateralized representation of gut cues in the brain in both mouse and humans^99^, and right-biased populations of putative gut-innervating reward neurons have been identified^35,100^. Stimulation of the vagus nerve is used in humans as treatment for a variety of diseases, including depression and epilepsy^101^. However, severe cardiac side effects sometimes result specifically from right-sided stimulation^102,103^. This asymmetric side-effect has been explained by the presence of both anatomical and functional asymmetries in cardiac innervation by the vagus nerve^104^. The asymmetry we identified in the *D. cerebrum* vagus is thus consistent with an evolutionarily broad trend in vagal organization.

While the role of the additional *piezo2b+/p2rx3b+* neurons on the right side of the sensory vagus is yet unknown, multiple sexual dimorphisms have been described at the anatomical and behavioral level in *D. cerebrum* adults. During male sexual maturation, the male-specific drumming apparatus develops from skeletal changes in association with the swim bladder, and the vent shifts anteriorly relative to the pelvic fin, concomitant with a rerouting of the digestive tract^105^. Adult sexual dimorphisms have also been identified in whole-brain activity patterns in response to male-specific vocalizations^106^. At the anatomical level, there are thus clear differences between adult male and female animals in the internal organs, brain, and body plan, which do not exist before sexual maturation. It has been repeatedly demonstrated in a variety of animals that neuron identity, neurotransmitter usage, and connectivity can differ significantly between the sexes in animals, and that these differences can drive sexually dimorphic behaviors^107–110^. Because male-specific vocalization in *D. cerebrum* is generated by mechanical drumming, it is tempting to speculate that the extra mechanosensory neurons provide some sort of sensory feedback to regulate this behavior. More broadly, our identification of molecular subtypes and anatomical organization in the *D. cerebrum* sensory vagus provides the foundation to link particular visceral neurons and sensory cues with their related behavioral outputs in both sexes.

## Acknowledgements

We thank the Judkewitz lab for generously providing *D. cerebrum* embryos to found our colony, and Alba Aparicio Fernandez, Rita Gonzalez, and Diana Medeiros Gomes for their expert care of the colony. Technical support for imaging was provided by the Biozentrum Imaging Core Facility (IMCF), and cell sorting was made possible by the Biozentrum FACS Core Facility. The Genomics Facility Basel, especially Christian Beisel and Mirjam Judith Feldkamp, supported library preparation and sequencing. Joaquín Navajas Acedo provided unpublished protocols and support for cell dissociation and FLASH-seq analysis. Vincent Hahaut and Simone Picelli granted access to an unpublished iteration of the FLASH-seq protocol^111^. We are grateful to Oded Mayseless, Will Joo, and Claude Desplan for critical reading of the manuscript. EAB received funding from the Jane Coffin Childs Fund for Medical Research and the University of Basel Fund for Excellent Junior Researchers.

## METHODS

### Fish Husbandry

A *Danionella cerebrum* colony was maintained under standard husbandry conditions for *Danio rerio*: 14-10h light-dark conditions, and a standard diet of Artemia twice per day and dry food once per day. All experiments were performed according to the Swiss Law and Kantonales Veterinäramt of Kanton Basel-Stadt (licenses #1035H and #3097).

Embryos were obtained from group breeding in standard holding tanks supplied with short lengths of silicone tubing, which serve as breeding shelters^21^.

### *p2rx3b:gfp* transgenic fish

To generate the *Tg(p2rx3b:gfp)* transgenic fish, we engineered a Tol2-GFP plasmid based on the previously reported *p2rx3.2::eGFP^GR^* plasmid^30^. We amplified 9kb upstream of the zebrafish *p2rx3b* start codon, the first exon, and part of the first intron (for a total size of 9.2kb) from genomic DNA and used NEBuilder HiFi DNA assembly to create Tol2-*p2rx3b*-GFP-Tol2. This was co-injected with Tol2 transposase mRNA. Larvae were screened at 3dpf for GFP expression and then raised to adulthood.

Although *D. cerebrum* breeds in groups, we were able to screen for individual F0 founders by crossing a single GFP+ F0 with a wild-type group of the opposite sex. Stable F1 transgenic progeny were raised to adulthood and outcrossed again to generate the transgenic line. All experiments were conducted on F2 and subsequent generations.

### HCR RNA *in situ* and immunohistochemistry

Following euthanasia in MS-222/Tricaine Methanesulfonate, fish were fixed overnight in 4% paraformaldehyde, and then dehydrated in a methanol series. Dehydrated fish were stored at −20C in 100% methanol at least overnight, and up to 6 months. For staining, fish were rehydrated in a decreasing methanol series (75%, 50%, 25% in phosphate buffered saline with 0.1% Tween). Pigment was removed using a 25 minute incubation in 0.8% potassium hydroxide and 0.9% hydrogen peroxide, also in 0.1% PBSTween.

Following washes in 0.1% PBSTween, animals were treated with 30ug/mL proteinase K for 45 minutes, then post-fixed for 20 minutes at room temperature in 4% PFA. Detection and amplification protocols for HCR RNA *in situ* followed an established protocol (Molecular Instruments, MI-Protocol-RNAFISH-Zebrafish, Revision Number 10). RNA probe sequences used are listed in **Supplemental Table 1**. In samples that were also stained using antibodies (Rabbit polyclonal SOX10, Lucerna Chem GTX128374 at 1:200, Chicken anti-GFP Invitrogen A10262 at 1:1000), primary antibody was added alongside HCR amplifiers, and secondary antibody incubation was performed overnight at 4C in 5X SSCT buffer. Secondary antibodies were all used at 1:500, and were ThermoFisher A10036 (donkey anti-mouse 546), ThermoFisher A11039 (goat anti-chicken 488), Jackson ImmunoResearch 703-545-155 (donkey anti-chicken 488),ThermoFisher A21235 (goat anti-mouse 647).

### Microscopy

Before imaging, all stained animals were incubated at least overnight in Ce3D Tissue Clearing Solution (BioLegend) at 4C. Fish were mounted on slides using silicone spacers between the slide and coverslip, which was flush with the cranial sensory ganglia (or brain where relevant). Point-scanning (LSM880, LSM980, Leica Stellaris) and spinning disc (Olympus SpinSR) confocal microscopy was used to image HCR RNA *in situ* and immunohistochemistry. For *p2rx3b* neuron tracing experiments animals were imaged using 10X air objectives, for HCR RNA *in situ* animals were imaged using immersion objectives (20X, 25X, or 30X). Multidimensional data was reconstructed as maximum intensity projections using Zeiss Zen, FIJI, or Imaris (Oxford Instruments) software.

For cell counting in HCR images, Z-stacks were viewed as volumetric projections in Imaris. The reference channel (*phox2bb* or *p2rx3b*) was used to create a mask defining the visceral vagus ganglion. The cell segmentation tool was used to mask cells with HCR puncta. Because there was not an independent measure of cell boundary, we consider masked cells to be an “arbitrary unit” of cell count. Segmentation parameters (expected cell size, seed point size) were kept consistent for each gene, so cell counts for a particular gene can be compared across animals.

For counting the number of neurons in p2rx3b:gfp images, the multipoint tool in FIJI was used to manually annotate each cell body. Z-stacks were quantified slice by slice to resolve overlapping neurons.

Traced p2rx3b:gfp neurites are displayed using the Imaris (Oxford Instruments) filament AutoPath tool.

### Tissue dissociation and cell sorting

For each sort, 12 animals were used. For each sex, sorts were performed on two separate days (24 total animals per sex). Adult *D. cerebrum* (2-4 months post fertilization) were euthanized in MS-222/Tricaine Methanesulfonate, and then immediately transferred to Dulbecco’s Phosphate Buffered Saline (DPBS) on ice. Animals were manually dissected using forceps to isolate the cranial sensory ganglia by removing the trunk, gut, heart, brain, jaw, and eyes. The dissected region was then transferred to dissociation buffer (9.45mg/mL protease from *Bacillus licheniformis* Sigma P5380, 0.5mg/mL DNaseI Sigma 10104159001, 2.5mM EDTA, 1.25X Gibco B-27 Supplement, all in DPBS) in a protein lo-bind Eppendorf tube. Every 3 minutes, the solution was mechanically disrupted by pipetting. After 24 minutes of dissociation (which was optimized via trials with cell sorting), the dissociation was stopped with the addition of 30% fetal bovine serum and 0.8mM CaCl2 in DPBS. DAPI (Sigma D9542) was added to identify dead cells. The solution was filtered through a 70 micron strainer, and diluted in resuspension buffer (5% fetal bovine serum, 1X Gibco B-27 Supplement, 0.8mM CaCl2 in DPBS) based on detected FACS concentration.

Sorts were performed using a BD FACSAriaIII with neutral density filter 1.5, gating for cell size (FSC and SSC), GFP expression, and live cells (DAPI). Sorts were performed at 4C. Cells were sorted directly into 384 well plates containing lysis buffer. At the end of each sort, plates were briefly centrifuged at 4C then flash-frozen on dry ice and transferred to −80C until library preparation. We were able to sort and sequence 873 cells (482 from males and 391 from females).

### Library preparation and sequencing

Single-cell RNA-seq libraries were prepared using the FLASH-seq protocol as previously described^32,111^ following the associated protocols.io (https://www.protocols.io/view/flash-seq-protocol-kxygxzkrwv8j). The I.DOT (Dispendix) was used for all dispensing steps, the Agilent Bravo system with 96ST head (Agilent Technologies) and Alpaqua 384 Post Magnet (NimaGen) for magnetic bead clean-up (SPRIselect reagent kit, Beckman Coulter) and the Mosquito HV system (SPT Labtech) for cDNA normalisation and setting up adapter ligation reactions. The RT-PCR reaction was performed with 21 cycles. QC of a subset of the cDNAs and the final library pools was done with the Fragment Analyzer HS NGS 1-6000 bp Kit (Agilent Technologies).

Concentrations were measured with Quant-iT PicoGreen (Thermo Fisher Scientific) using the Infinite M1000 Pro (Tecan). The tagmentation reaction was set up with “inhouse” Tn5 transposase^112^ (produced by the Protein Production and Structure Core Facility, EPFL). Indexing primers were ordered from IDT; sequences of primers are available on protocols.io. Library pools were sequenced SR76 with NextSeq 500/550 High Output Kit v2.5 (75 Cycles) (Illumina).

### FLASH-seq data analysis

For initial read alignment and filtering, we used the STAR software pipeline^113^. For each cell, paired-end read FASTQ files were aligned together to the *D. cerebrum* reference genome (GenBank assembly GCA_007224835.1^114^) to identify uniquely-mapped reads and filter out multi-mapping or unmapped reads. The subread package (https://github.com/ShiLab-Bioinformatics/subread) was then used to generate a feature counts matrix from the mapped reads. This counts matrix was loaded into R and used to generate a Seurat object of the dataset.

After filtering out cells with low numbers of features (<200) or RNA counts (<10,000), and high numbers of features (>12,000, suggesting two cells could have been sorted into the same well), 803 of our 873 sequenced cells remained and were used for downstream analysis. We found that the in-plate library preparation and sequencing approach utilized by FLASH-seq introduced some bias in sequencing depth (cells along the borders of each 384 well plate were generally sequenced to a higher depth than cells in the center of the plate). For this reason, we used the Seurat scTransform pipeline, which was specifically developed to address this type of technical heterogeneity, to normalize the data^33,115^. We performed dimensionality reduction using 30 PCs; as reported, we found that the results were robust to this parameter when using scTransform. We used the Wilcoxon rank-sum test (FindAllMarkers) to identify differentially expressed genes between the clusters.

We used the SAMap package^116^(https://github.com/atarashansky/SAMap/tree/main) to perform BLAST-based mapping between the *D. cerebrum* genome and the *Danio rerio* GRCz11 genome assembly. We filtered this “dictionary” to generate a list containing the highest-ranked zebrafish ortholog (by E value) for each *D. cerebrum* gene. We used this dictionary only for interpretation of our marker genes: all clustering and statistical analysis were performed using the native *D. cerebrum* gene annotations, and genes were not filtered or excluded based on whether a high confidence *D. rerio* ortholog was identified. The first time we reference any gene name throughout the text, we provide both the zebrafish common name and the annotated *D. cerebrum* gene name. In subsequent references we simply use the zebrafish common name.

For assessing conservation with the mouse sensory vagus, we manually curated a list of marker genes identified across multiple studies^1,6,34^. We performed protein BLAST against the *D. cerebrum* reference genome to identify a permissive (one-to-many) ortholog list for each mouse marker gene. This ortholog list was then intersected against the marker gene list to identify which mouse vagus marker orthologs were also statistically significant marker genes for our cranial ganglion clusters.

For transcripts that were used to design HCR probes, we also used our paired end sequencing to verify the reference genome transcript annotation using the StringTie package^117^. In some cases, this suggested errors in the original annotations. These were most often regarding exon exclusion or inclusion within correctly annotated gene boundaries, but in other cases involved annotation of gene boundaries. For instance, in the case of *piezo2a/*DNTS_004436/DNTS_004431, we believe that two neighboring annotations (DNTS_004436 and DNTS_004431) actually encode a single gene orthologous to zebrafish *piezo2a*.

### Figure preparation

Plots were generated in R using the Seurat, beeswarm, and ggplot2 packages. Statistical tests as indicated in the figure legends were performed in R. Figures were prepared using Adobe Photoshop and Adobe Illustrator.

## Notes

### Competing Interest Statement

A.F.S. is a scientific advisor to Novartis.

