## Supplemental Figures and Legends for "Transcriptional subtypes, anatomical organization, and sexual dimorphism of sensory vagus neurons in *Danionella cerebrum*"

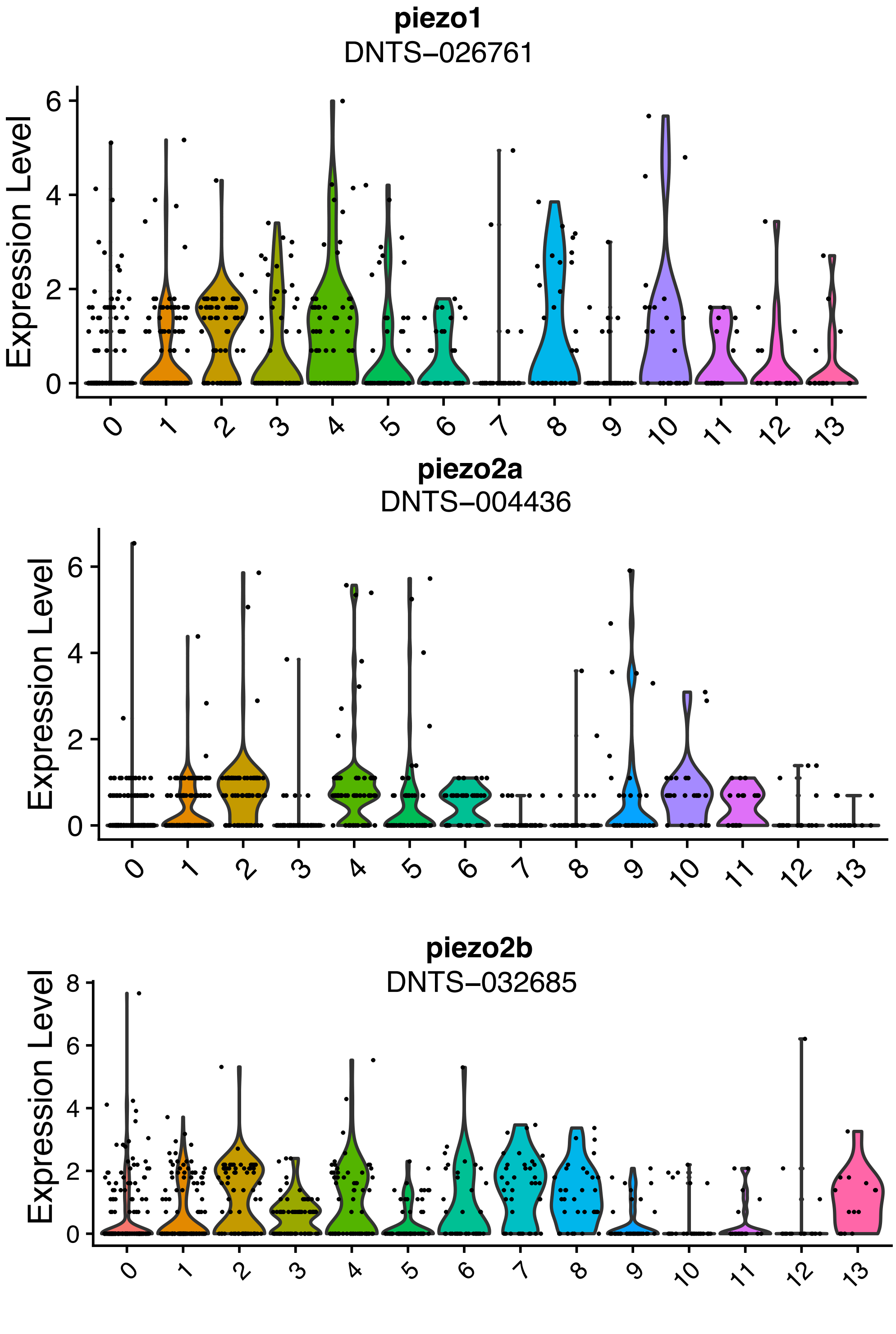


**Supplemental Figure 1**

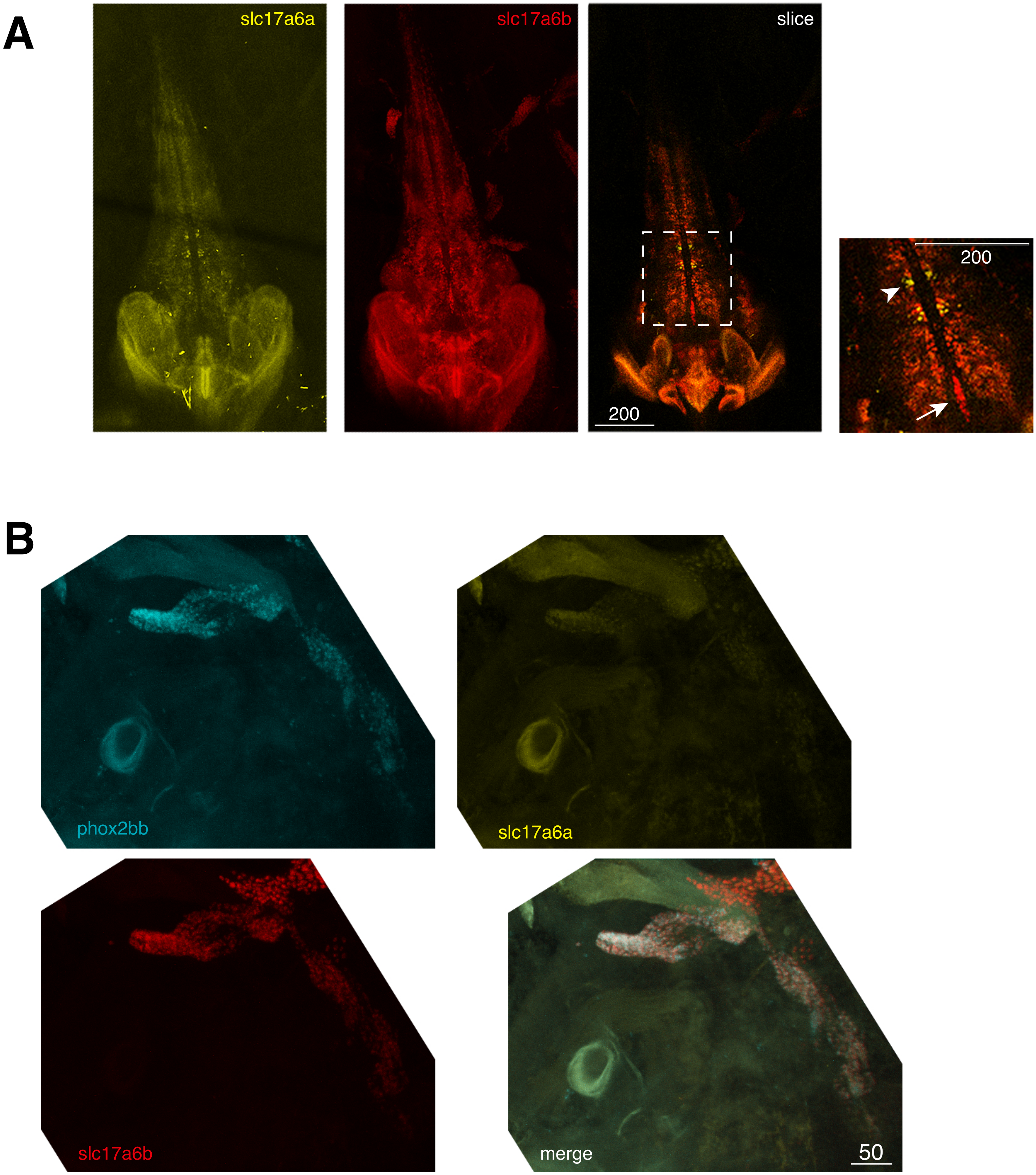


**Supplemental Figure 2**

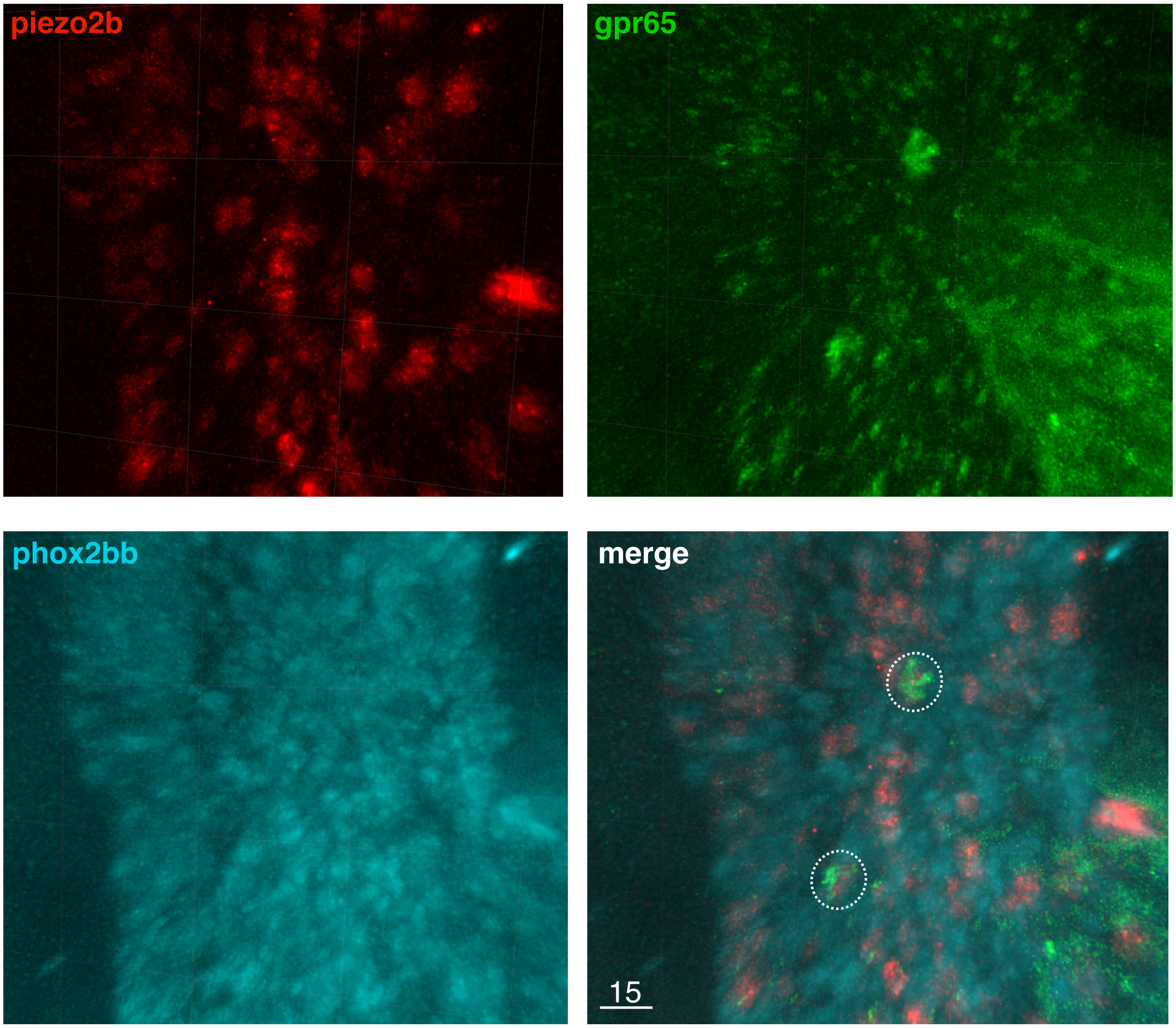


**Supplemental Figure 3**

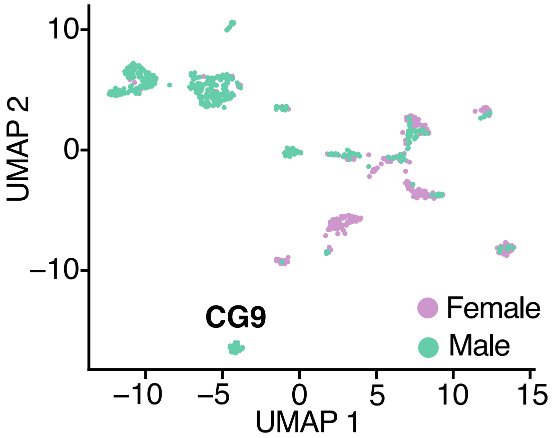


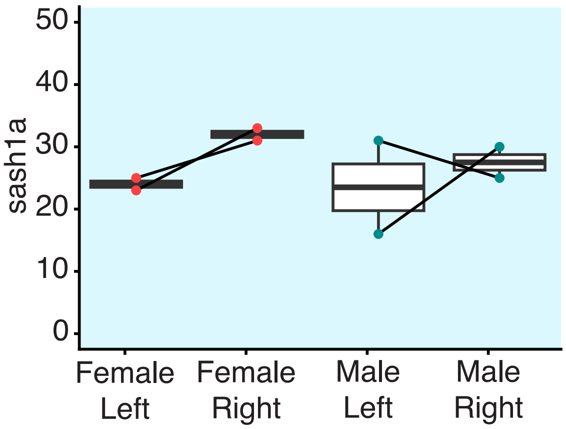


**Supplemental Figure 4**

**Supplemental Figure 1: Piezo mechanosensory channels are broadly expressed in p2rx3b:gfp+ subclusters.** Violin plots showing SCT-normalized expression of *piezo1*, *piezo2a*, and *piezo2b* across the 14 transcriptional subclusters.

**Supplemental Figure 2: Vglut2 paralogs are differentially expressed in the sensory vagus and central nervous system.**

**A: *slc17a6a* and *slc17a6b* expression in the adult brain.** *slc17a6a* and *slc17a6b* are expressed in overlapping and non-overlapping populations of neurons. Orthogonal projection showing a whole adult brain from the dorsal side (anterior down), with *slc17a6a* on the left, *slc17a6b* center, and a merged single slice image on the right to demonstrate non-overlapping expression. White dashed inset is shown magnified to the right. Arrowhead indicates neurons exclusively expressing *slc17a6a,* and arrow indicates neurons exclusively expressing *slc17a6b*. White scale bars shown with scale in microns in both panels.

**B: *slc17a6a* and *slc17a6b* expression in the sensory vagus**. *slc17a6a,* but not *slc17a6b*, is expressed in the adult sensory vagus. Orthogonal projections showing each channel individually, with *phox2bb* for anatomical reference, with channels merged in the bottom right.

**Supplemental Figure 3: The sensory vagus contains polymodal neuron types.** Orthogonal projection of HCR RNA *in situ* staining of *piezo2b, gpr65,* and *phox2bb*. Merged image of all three channels is shown in bottom left, with co-expressing cells encircled in white dashed circles. The *gpr65* images are a higher zoom inset from the same animal as **Figure 6D**, with the *piezo2b* channel added to indicate colocalization of *piezo2b* with the high *gpr65* cells.

**Supplemental Figure 4: The sensory vagus does not contain a male-specific molecular subtype.**

One of our scRNAseq clusters, CG9, contained only cells from male animals. However, we found by HCR for the *sash1a*/DNTS_007915 marker that an equal number of cells were present in both males and females (and on the left and right sides), suggesting that this was just a technical sampling bias in our sequencing. Left, UMAP representation from Figure 2A showing male vs. female identities of cells, with CG9 indicated. Right, quantification of *sash1a* HCR RNA *in situ* signal in adult males and females on the left and right sides.

**Supplemental Table 1: Sequences of DNA probes used for HCR RNA *in situ*.**
